# A Systematic Benchmark of Antibiotic Resistance Gene Detection Tools for Shotgun Metagenomic Datasets

**DOI:** 10.64898/2026.02.04.703716

**Authors:** Sumeet K. Tiwari, Alise J. Ponsero, Samuel Haynes, Judit Tálas, Kal P. Grimes, Andrea Telatin

## Abstract

Accurate detection of antimicrobial resistance genes (ARGs) from metagenomic data is essential for understanding resistance dissemination within microbial communities, yet tool performance remains influenced by sequencing coverage, community complexity, and dataset variability. In this study, we systematically benchmarked five widely used read-based ARG detection tools (ARGprofiler, KARGA, ARIBA, GROOT, and SRST2) across simulated metagenomic datasets representing varying sequencing coverages, microbial complexities, and approximate realistic metagenomic dataset. The results demonstrated that sequencing coverage is a major determinant of ARG detection accuracy, with reliable detection achieved at 10× coverage and performance stabilizing between 20× and 30×. ARGprofiler exhibited the highest overall F1-score (0.891 at ≥10×), whereas KARGA showed higher recall at low coverage levels, but lower precision compared to ARGprofiler. Increasing community complexity led to a decline in accuracy across all tools, and under realistic uneven coverage, performance variability increased substantially, with KARGA achieving the highest mean F1- score (0.122 ± 0.067). Runtime evaluation further revealed substantial differences in computational efficiency, with ARGprofiler, SRST2, and GROOT being the most resource-efficient, while KARGA imposed the highest computational burden. Collectively, these findings highlight that both sequencing coverage and community complexity profoundly shape ARG detection outcomes and that tool selection should balance accuracy with computational efficiency. The study also emphasizes the need for standardized benchmarking datasets that reflect true metagenomic complexity to ensure robust and comparable ARG surveillance across analytical pipelines.

## 2. Introduction

Antimicrobial resistance (AMR) is a major global health crisis, responsible for an estimated 1.27 million deaths in 2019 and projected to rise to nearly 2 million annually by 2050 ^1,2^. The spread of antimicrobial resistance genes (ARGs) across microbial communities threatens antibiotic effectiveness and underscores the urgent need to understand their distribution and dynamics for effective surveillance and interventions ^1,2^. Recent advancements in sequencing technologies, particularly shotgun metagenomic sequencing, have transformed the study of microbial communities by enabling culture-independent taxonomic and functional profiling directly from environmental samples ^3–5^.This approach allows the simultaneous detection of functionally relevant genes, such as antimicrobial resistance determinants, across hundreds of species ^6,7^ .

Despite these advances, the reliable detection and quantification of ARGs from metagenomic data remain challenging. The inherent complexity of microbial communities, including high species diversity, and closely related gene variants complicate identification of ARGs. At the sequencing level, factors such as variable depth, uneven coverage, and fragmented nature of metagenomic assemblies hinder the detection of ARGs ^8,9^. ARG detection methods employ diverse algorithmic approaches, each with distinct performance trade-offs. These ARG detection tools can be broadly characterized into following groups: (1) **Read-mapping based tools**, which directly align metagenomic reads against curated ARG databases for rapid profiling^10–12^; (2) **Assembly-based**, which requires contigs or genomes to enable detection of both known and novel ARGs^13–15^; but may face challenges with low-abundance genes; and (3) **Machine learning and deep learning methods**, which leverage sequence features, hidden Markov models, or neural networks to enhance sensitivity, particularly for novel or highly divergent ARGs^16–19^. Each tool varies substantially in their precision, recall, computational requirements, and suitability for different types of datasets. Each tool required a reference database in a dedicated format.

Similarly, ARGs databases differ in their content and curation strategies. The Comprehensive Antibiotic Resistance Database (CARD) prioritizes experimentally validated, clinically relevant determinants and provides detailed ontology-based annotations^20,21^. ResFinder targets characterized ARGs primarily from clinical isolates^22^. MEGARES and BacMet extend coverage to include metal and biocide resistance genes^23,24^, while functional metagenomics-based database like ResFinderFG captures the ARGs discovered from environmental samples^25^.

A benchmarking study has evaluated the performance of ARG detection tools in the context of metagenomic samples using different databases^26^. The study has concluded that different combinations of tools and reference databases produce significantly different results^26^. This variability likely results from distinct coverage and annotation depth of ARGs in the databases, uneven representation of ARG subtypes, the inherent biological complexity of metagenomic samples, and whereas another study has shown the impact of sequencing depth and coverage^27^.

Systematic comparisons of ARG detection tools on metagenomic data are limited. Most benchmarking efforts have focused on isolate-based detection, which involves reduced biological complexity compared to metagenomic samples. Consequently, the influence of key factors such as microbial diversity, sequencing depth, and uneven coverage on the performance of ARG detection tools in metagenomic settings remains insufficiently explored. Therefore, to begin with, we evaluated a subset of read-level ARG detection methods that operate independently of metagenome assembly and can leverage a harmonized ARG database, thereby controlling for differences in representation of ARGs, and their subtypes.

Specifically, we comprehensively benchmarked widely used read-level ARG detection tools using simulated shotgun reads from mock-communities. The simulated dataset enabled us to systematically vary sequencing coverage and biological complexity. Our objectives were to assess the impact of sequencing coverage and microbial complexity on detection accuracy, given a harmonized ARG database as well as to evaluate computational efficiency and resource requirement for each tool. All the tools were executed in Singularity^28^ containers via Nextflow^29^ pipeline to ensure reproducibility.

Unlike previous benchmarking studies, which often rely on isolate genomes or fixed sequencing conditions, this work systematically disentangles the effects of sequencing coverage, microbial community complexity, and uneven coverage distributions using harmonized reference databases. This design enables a more realistic and controlled assessment of ARG detection performance in metagenomic contexts.

## 3. Material and Methods

### a. Database Preparation

The bacterial genes conferring resistance to antibiotics, and biocides were collected from publicly available resources. Becauseheavy metals are known to co-select for antibiotic resistance ^30^, genes associated with metal resistance were also included. The complete database preparation workflow is illustrated in Figure 1. To create a comprehensive resistance genes collection, the following databases were utilized: AMRFinderPlus (v3.12)^14^, ARG-ANNOT (v6, July 2019)^31^, BacMet (v1.1, https://doi.org/10.5281/zenodo.8108201)^32,33^, CARD (v3.3.0)^21^, MegaRes (v3.0)^24^, ResFinderFG (v2.0) and ResFinder^22,25^.

**Figure 1:**
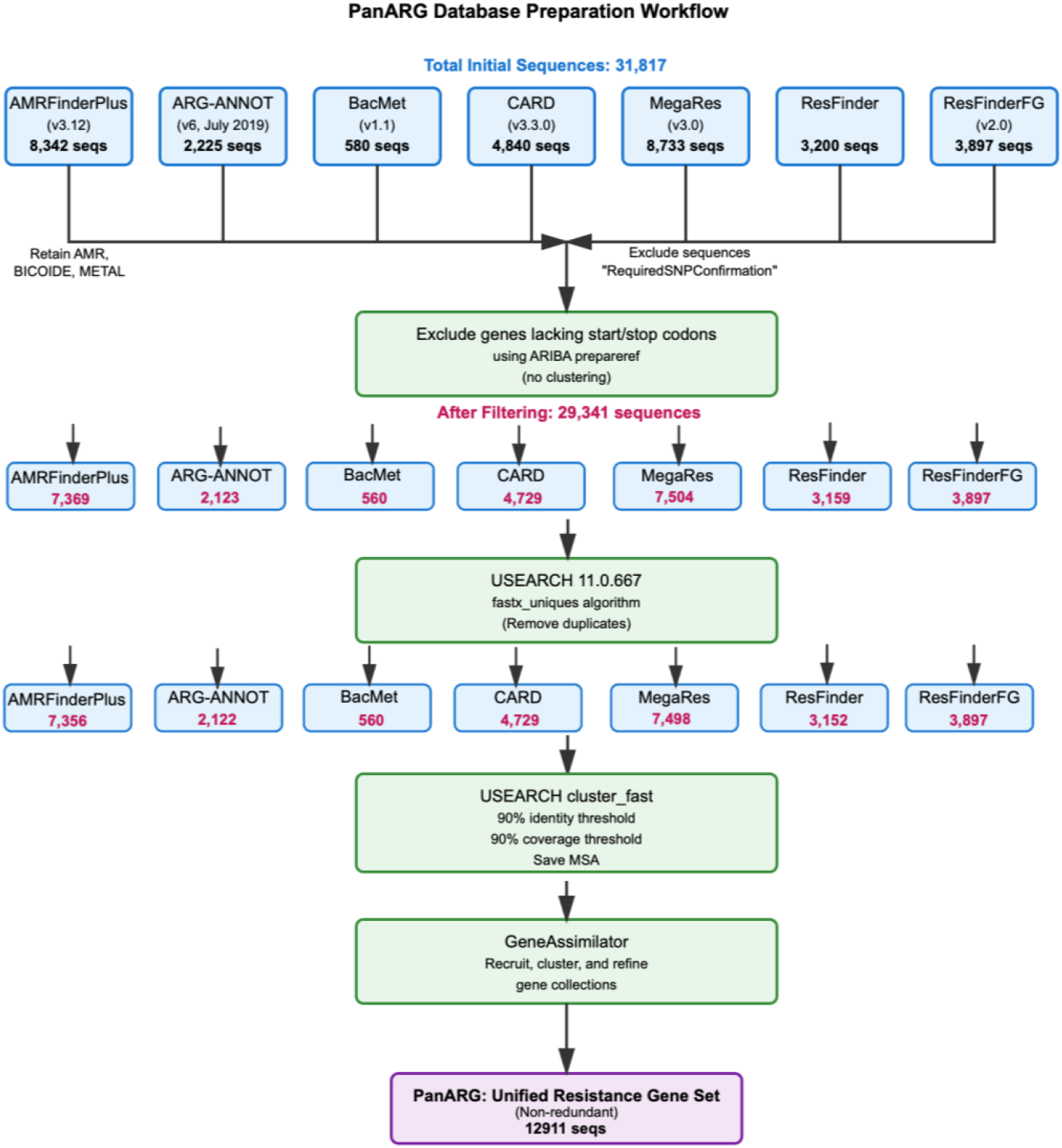
Workflow for resistance gene database compilation and curation: Showing the collection and processing of antimicrobial, biocide, and metal resistance genes from seven public databases (AMRFinderPlus, ARG-ANNOT, BacMet, CARD, MegaRes, ResFinder, and ResFinderFG). Initial sequences (n=31,817) were filtered to remove genes lacking start/stop codons (n=29,341 retained), deduplicated, clustered at 90% identity/coverage, and refined using GeneAssimilator to generate a unified resistance gene collection.

From the CARD database, the genes based on protein homolog model were included as they do not rely on mutation for resistance. In MegaRes gene collection, sequences with the “RequiredSNPConfirmation” tag were excluded, as these genes required SNPs to confer resistance. From the AMRFiderPlus database, sequences categorised under the subtype “AMR”, “BIOCIDE” and “METAL” were retained. Additionally, genes which lack start and stop codons were also excluded from all the collected databases using the “ARIBA prepareref” module ^34^, without clustering the sequences, as such genes might be non-coding, partial or truncated genes.

Unique sequences were then identified by clustering using USEARCH 11.0.667 with the fastx_uniques algorithm ^35^. These unique sequences were clustered based on 90% identity and 90% coverage^33^ using the cluster_fast algorithm and the multiple sequence alignments (MSA) were saved. GeneAssimilator (https://github.com/genomicepidemiology/gene_assimilator) was used to perform this iterative approach to recruit, cluster, and refine the gene collections of various sources into a unified set.

### b. Preparation of simulated dataset

An overview of the complete benchmarking is represented in Figure2. A mock microbial community was built using two publicly available datasets. The first dataset is the HumGut genome collection ^36^ which is a comprehensive repository of bacterial genomes representative of healthy human gut microbiota. The second dataset comprises a collection of highly resistant clinical bacterial isolates^26^ . The microbial genomes were screened for the presence of ARGs using ABRICATE^13^ (with default parameters) against the unified panARG database. These genomes were then divided into two groups: a resistant group (having at least one predicted ARG) and a non-resistant group (with no ARG; Figure 2a). Three experimental designs were implemented to evaluate ARG detection performance (Figure 2b):

i. **Effect of sequencing coverage:** To investigate how coverage influences the detection of antimicrobial resistance genes (ARGs), the microbial composition was kept constant. A mock community of 20 bacterial genomes (Table S1) was therefore constructed, comprising 10 genomes carrying known ARGs and 10 genomes without ARGs. Illumina HiSeq paired-end reads were simulated from the mock community using InSilicoSeq^37^ at 21 different coverage levels. The coverage levels included 1×, followed by increments of 5× from 5× to 100× (i.e., 1×, 5×, 10×, 15×, …, 100×). The simulated datasets were then analysed using ARG detection tools to evaluate how sequencing depth affects the sensitivity and accuracy of ARG gene identification.
ii. **Effect of microbial complexity:** To assess how microbial complexity impacts the detection of ARG genes, an initial set of 30 genomes (15 ARG-positive and 15 ARG-negative) was randomly selected (Table S2) Illumina HiSeq paired-end reads were simulated for this set using InSilicoSeq^37^ at 25× sequencing depth. The dataset was expanded iteratively by adding 10 genomes (5 ARG-positive and 5 ARG-negative) in each iteration, up to a total of 100 iterations. For each iteration, reads were simulated and analysed using ARG gene detection tools. Sequencing depth was fixed at 25×, based on the results of the coverage analysis, to ensure that any observed differences could be attributed specifically to microbial diversity rather than variation in coverage.
iii. **Real mock-community:** To approximate a realistic metagenomic community, dRep (v3.4.2)^38^ was used dereplicate HumGut genomes meeting quality thresholds of ≥90% completeness and ≤5% contamination was retained (n = 2,640 of 30,691). From this dereplicated set, 100 genomes representing 100 distinct species were randomly selected to construct 100 independent mock communities. Illumina HiSeq reads were then simulated for each community with a lognormal coverage distribution in InSilicoSeq.

**Figure 2:**
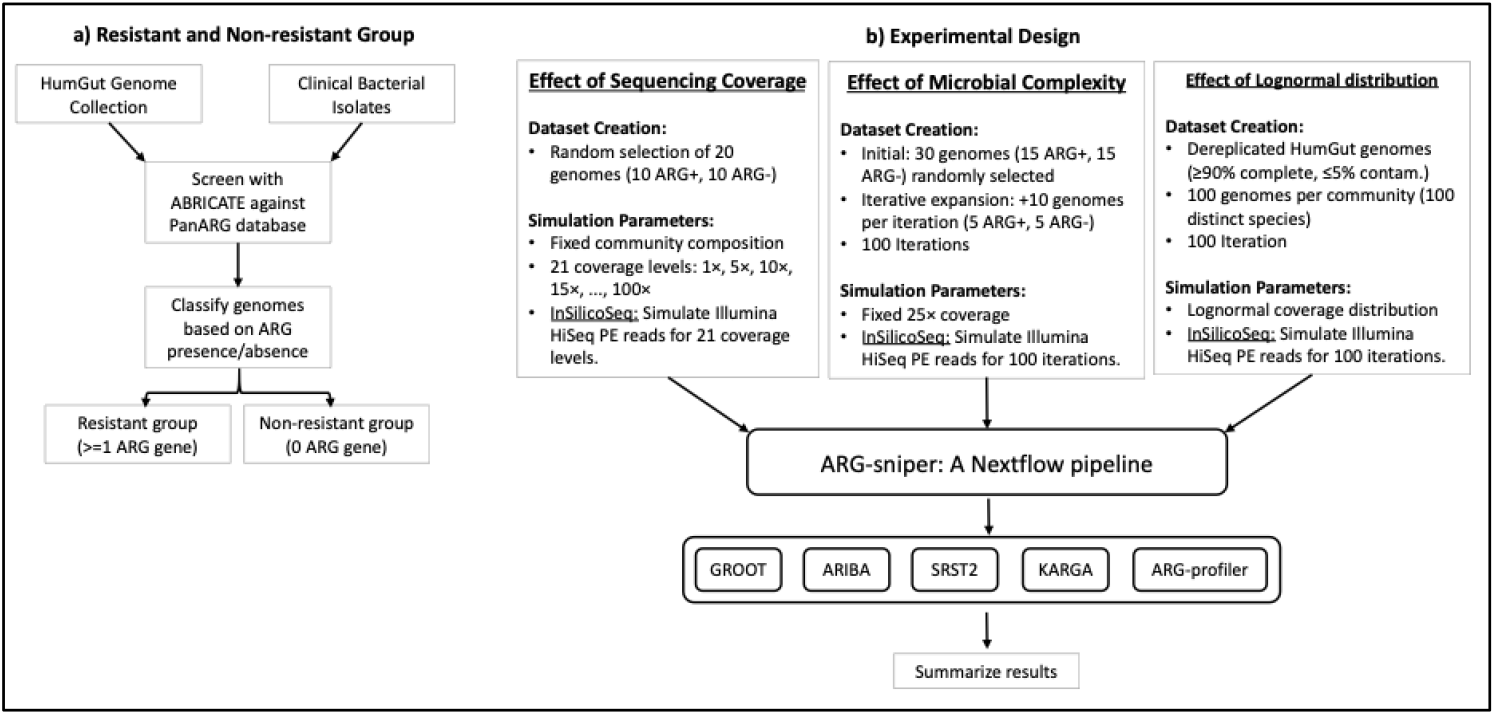
Workflow for benchmarking ARG detection tools. (a) Resistant and non-resistant group. Mock microbial communities were constructed from HumGut genome collection and clinical bacterial isolates. Genomes were screened for ARGs using ABRICATE against the PanARG database and divided into resistant (≥1 ARG) and non-resistant (no ARG) groups. (b) Experimental Design and Analysis pipeline. Three mock community designs assessed tool performance under different coverage levels, microbial complexity, and lognormal distribution of coverage. Simulated reads were screened for ARGs via Nextflow pipeline with selected ARG detection tools.

### c. Criteria for selection of tools

To assess the performance of different reads-based ARG detection tools, we defined a set of selection criteria to ensure consistency, comparability, and practical utility. We established the following criteria: a) the input should be FASTQ files rather than assemblies. b) the tool must allow the use of a custom ARG database not protein database. c) the tool must be executable via the command line and d) only local assembly is permitted. Based on these criteria, five tools were selected: GROOT^39^, ARIBA^34^, SRST2^40,41^, KARGA^41^ and ARG-profiler^33^. A custom Nextflow (v22.10.7) pipeline was developed incorporating these tools as modules in the pipeline (Figure 2b). The list of modules as follows: a)GROOT b) ARIBA c) SRST2 d) KARGA e) ARGprofiler (only ARG detection module using KMA). However, both ARIBA and SRST2 were original developed to predict ARGs from isolates, not for metagenome.

The performance of these tools was compared against the ARGs detected by ABRICATE, an assembly-based method that uses BLAST for gene identification. Assembly-based detection was chosen as the comparison baseline because it eliminates read-mapping ambiguities and enables more complete gene recovery from assembled contig. Although assembly-based approaches may miss low-abundance or fragmented genes, ABRICATE provides a conservative and reproducible reference for benchmarking read-based methods when applied with stringent identity and coverage thresholds. Assemblies used in the simulating reads for each experimental design were screened with ABRICATE against the panARG database using 90% identity and query coverage thresholds.

## 4. Results

### a. Unified Resistance Gene Database

A comprehensive resistance gene database was compiled from seven publicly available resources, yielding an initial collection of 31,817 sequences (Figure 1). These included genes conferring resistance to antibiotics, biocides, and heavy metals from AMRFinderPlus (n=8,342), ARG-ANNOT (n=2,225), BacMet (n=580), CARD (n=4,840), MegaRes (n=8,733), ResFinder (n=3,200), and ResFinderFG (n=3,897). A total of 2,476 (7.8%) sequences that lacked start and/or stop codons were excluded, resulting in 29,341 high-quality sequences. Subsequent deduplication and clustering at 90% identity and coverage thresholds using USEARCH, followed by iterative refinement with GeneAssimilator,produced a unified, non-redundant resistance gene collection named as panARG.

The comparative analysis of ARG gene databases revealed a substantial overlap in genes among the ARG databases (in Figure 3a). A total of 23,658 (80.6%) of genes were found to be shared among the databases, whereas 5,683 (19.4%) are unique to individual databases (in Figure 3a). None of the genes were shared across all seven databases (in Figure 3a). The pairwise Jaccard similarity analysis revealed the substantial heterogeneity in gene content across the seven ARG databases (in Figure 3b). Overall, the mean Jaccard similarity index was low (≈ 0.21), and the mean Jaccard distance between databases was 0.79. The highest concordance was observed between AMRFinderPlus and MEGARES (J=0.66), followed by CARD-MEGARES (J=0.59) and AMRFinderPlus-CARD (J=0.56). In contrast, ResFinderFG and BacMet exhibited least overlap with other databases (J < 0.01 and J < 0.07, respectively).Notably, ResFinderFG contained the highest number of unique genes, followed by AMRFinderPlus and MEGARES (Figure 3a).

**Figure 3:**
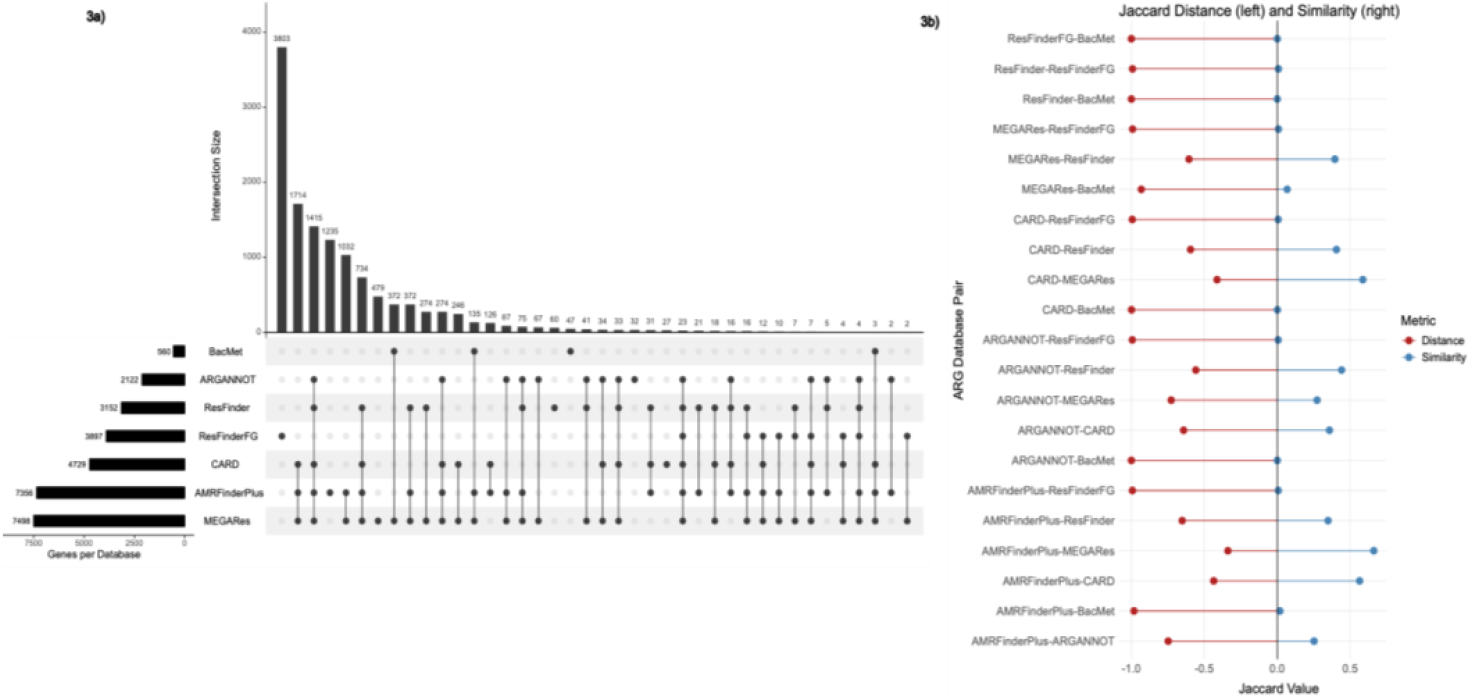
Comparative analysis of ARG databases. **(a)** UpSet plot showing the distribution and intersections of genes across seven ARG databases. Horizontal bars (left) indicate total genes per database. Vertical bars show intersection sizes, with filled circles and connecting lines indicating which databases contribute to each intersection. **(b)** Lollipop plot displaying pairwise Jaccard similarity and distance between ARG databases. Each database pair extends from the central axis, with similarity indicating gene overlap and distance representing dissimilarity (1 - similarity).

### b. Effect of sequencing coverage

We assessed five AMR detection tools (ARGprofiler, ARIBA, KARGA, SRST2, and GROOT) using simulated metagenomic datasets at 21 different sequencing depths ranging from 1× to 100× coverage. Performance was evaluated based on the detection of AMR genes, with abricate-based predictions serving as the reference, and assessed using recall, precision, F1 score and Coefficient of Variation (CV) (Table 1a-b, Figure 4a).

**Table 1a:** Tool performance across sequencing depth (1x-100x). CV = Coefficient of Variation (SD/Mean × 100) calculated for F1 scores only.

| Tool | F1-Score Mean ± SD | F1-Score Range | Precision Mean ± SD | Precision Range | Recall Mean ± SD | Recall Range | F1 CV (%) |
| --- | --- | --- | --- | --- | --- | --- | --- |
| ARGProfiler | 0.815 ± 0.236 | 0.000-0.903 | 0.861 ± 0.194 | 0.000-1.000 | 0.805 ± 0.246 | 0.000-0.909 | 29.0 |
| ARIBA | 0.724 ± 0.177 | 0.022-0.802 | 0.754 ± 0.060 | 0.500-0.817 | 0.729 ± 0.199 | 0.011-0.841 | 24.4 |
| KARGA | 0.522 ± 0.116 | 0.044-0.617 | 0.587 ± 0.166 | 0.416-1.000 | 0.527 ± 0.121 | 0.023-0.602 | 22.2 |
| SRST2 | 0.190 ± 0.124 | 0.000-0.395 | 0.176 ± 0.104 | 0.000-0.333 | 0.315 ± 0.259 | 0.000-0.705 | 65.4 |
| GROOT | 0.182 ± 0.057 | 0.000-0.233 | 0.267 ± 0.095 | 0.000-0.600 | 0.147 ± 0.052 | 0.000-0.193 | 31.3 |

**Table 1b:**
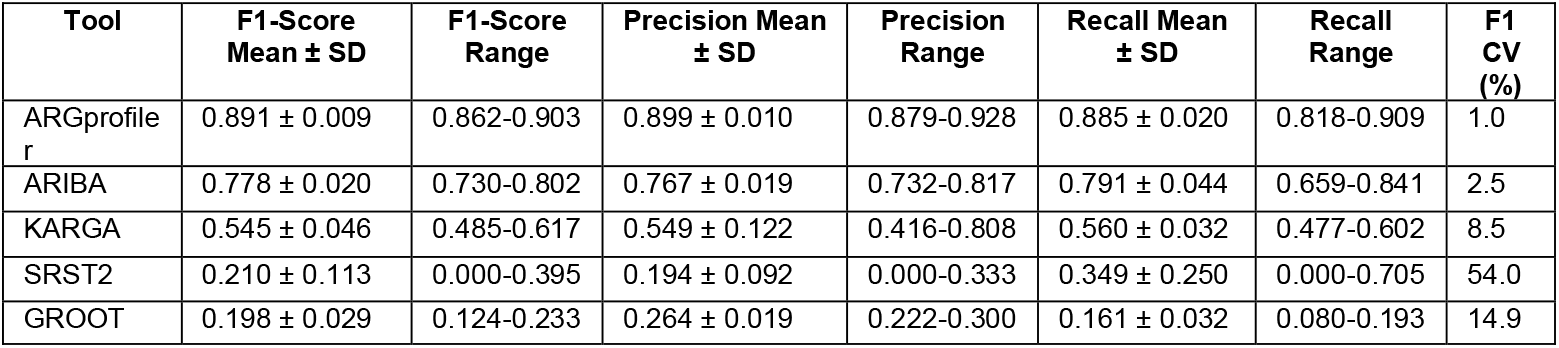
Tool Performance from 10x to 100x, CV = Coefficient of Variation (SD/Mean × 100) calculated for F1 scores only.

| Tool | F1-Score Mean ± SD | F1-Score Range | Precision Mean ± SD | Precision Range | Recall Mean ± SD | Recall Range | F1 CV (%) |
| --- | --- | --- | --- | --- | --- | --- | --- |
| ARGprofile r | 0.891 ± 0.009 | 0.862-0.903 | 0.899 ± 0.010 | 0.879-0.928 | 0.885 ± 0.020 | 0.818-0.909 | 1.0 |
| ARIBA | 0.778 ± 0.020 | 0.730-0.802 | 0.767 ± 0.019 | 0.732-0.817 | 0.791 ± 0.044 | 0.659-0.841 | 2.5 |
| KARGA | 0.545 ± 0.046 | 0.485-0.617 | 0.549 ± 0.122 | 0.416-0.808 | 0.560 ± 0.032 | 0.477-0.602 | 8.5 |
| SRST2 | 0.210 ± 0.113 | 0.000-0.395 | 0.194 ± 0.092 | 0.000-0.333 | 0.349 ± 0.250 | 0.000-0.705 | 54.0 |
| GROOT | 0.198 ± 0.029 | 0.124-0.233 | 0.264 ± 0.019 | 0.222-0.300 | 0.161 ± 0.032 | 0.080-0.193 | 14.9 |

**Figure 4:**
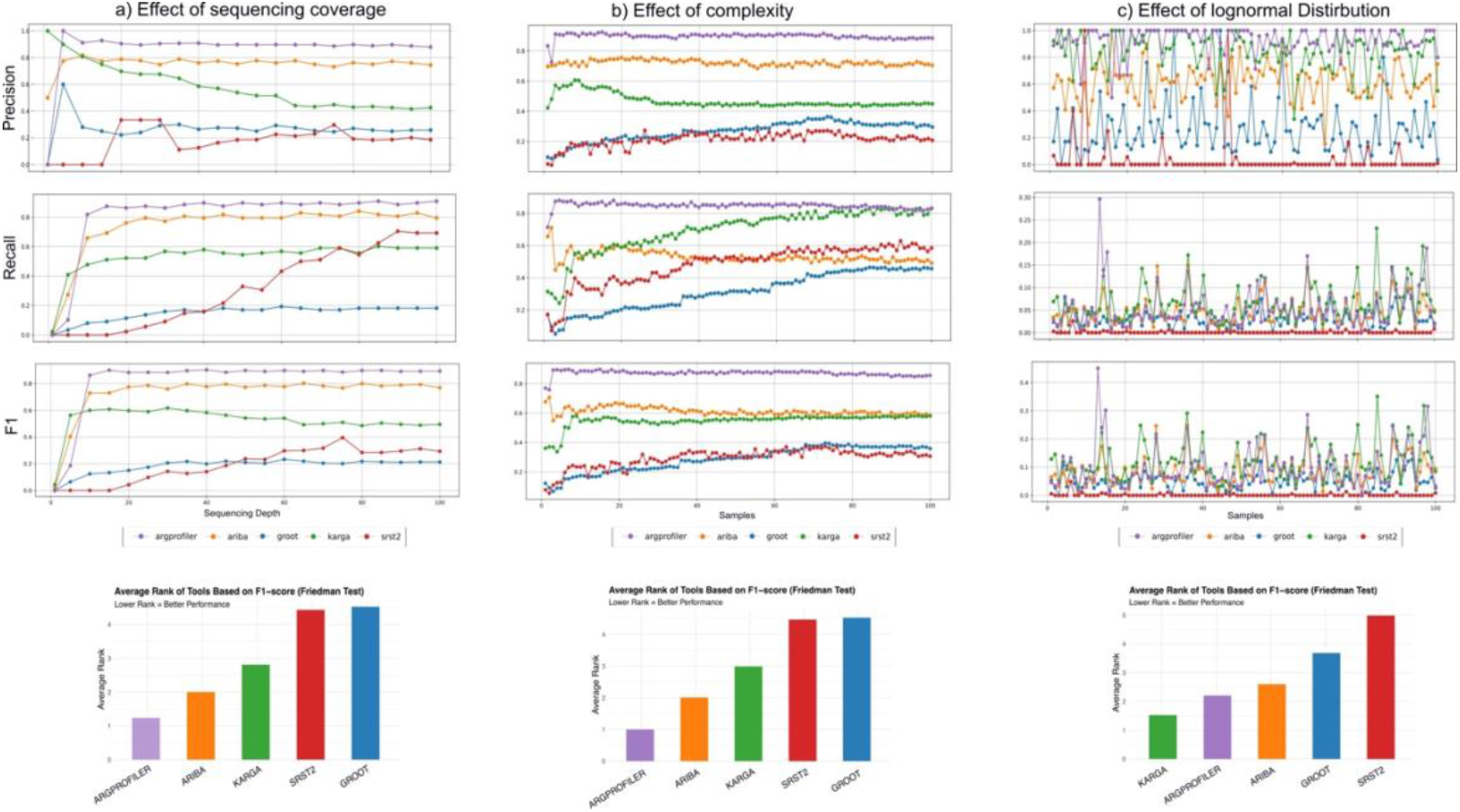
Performance of ARG detection tools across three experimental designs. Five ARG detection tools (ARGprofiler, ARIBA, KARGA, GROOT, and SRST2) was evaluated on simulated metagenomic datasets under three experimental conditions: **(a)** varying sequencing coverage **(b)** increased community complexity, and **(c)** log-normal coverage distribution mimicking realistic metagenome dataset. For each condition, tool performance was assessed using precision, recall and F1-score. The above panel indicate the evaluation metrices per sample in each experimental design. The bottom panel in each subplot shows the mean Friedman rank, based on F1-scores summarizing relative performance (left to right: higher to lower) of the tools under each condition.

ARGprofiler and ARIBA demonstrated high performance at variable sequencing depth. ARGprofiler achieved mean F1-score of 0.815 ± 0.236 (recall=0.805 ± 0.246; precision=0.861 ± 0.194; CV=29%), with performance stabilizing at >= 10x coverage (mean F1=0.891 ± 0.009; precision= CV=1%; see Table 1b). ARIBA yielded mean F1-score = 0.724 ± 0.177 (recall = 0.729 ± 0.199; precision = 0.754 ± 0.060; CV = 24.4%) improving at coverage above 10× (F1-score = 0.778 ± 0.020; CV = 2.5%).

KARGA showed moderate overall performance, yielding mean F1-score = 0.522 ± 0.116 (recall = 0.527 ± 0.121; precision = 0.587 ± 0.166; CV = 22.2%), with modest improvement at ≥10× coverage (F1-score = 0.545 ± 0.046; CV = 8.5%).

SRST2 and GROOT consistently performed poor, yielding F1-scores of 0.190 ± 0.124 and 0.182 ± 0.057, respectively, with correspondingly lower recall and precision values across all sequencing depths. SRST2 exhibited CV = 65.4%, while GROOT showed CV = 31.3%. At ≥10× coverage, both tools showed reduced variability (SRST2 CV = 54%; GROOT CV = 14.9%), though F1-scores remained below 0.3.

A Friedman rank-sum test revealed a significant difference in F1-scores among the tools (χ^2^= 72.12, df = 4, p = 8.11 × 10^−15^). Pairwise comparisons using the Nemenyi post-hoc test (single-step adjusted p-values) indicated that most tool pairs differed significantly (p < 0.05), except for ARIBA and ARGprofiler (p = 0.5223), KARGA and ARIBA (p = 0.4596), and GROOT and SRST2 (p = 0.9997). Significant differences were observed between GROOT and ARGprofiler (p = 1.7 × 10^−10^), GROOT and ARIBA (p = 2.3 × 10^−6^), KARGA and ARGprofiler (p = 0.0112), KARGA and GROOT (p = 0.0040), and SRST2 with all tools except GROOT (p < 0.01). The relative ranking of the tools based on their mean F1-scores is ARGprofiler > ARIBA > KARGA > SRST2 ≈ GROOT (Figure 4a).

### c. Effect of increase in complexity

The impact of sample complexity on these tools was assessed by using 100 systematically constructed mock communities. Complexity was progressively increased by adding more genomes. The assessment began with began with 30 genomes representing 28 unique species and 38 AMR genes and ended with 1,020 genomes representing 589 unique species and 846 AMR genes (Table S1). To disentangle the influence of sequencing coverage from community diversity, sequencing coverage was fixed at 25× across all datasets. with ABRICATE-based predictions serving as the reference, and assessed using recall, precision, F1 score and Coefficient of Variation (CV) (Table 2, Figure 4b).

**Table 2:** Comprehensive Tool Performance Metrics Across 100 Datasets *CV = Coefficient of Variation (SD/Mean × 100)

| Tool | F1 Score Mean ± SD | F1 Score Range | Precision Mean ± SD | Precision Range | Recall Mean ± SD | Recall Range | F1-Score CV (%) |
| --- | --- | --- | --- | --- | --- | --- | --- |
| ARGprofiler | $0.872 \pm 0.019$ | 0.759-0.897 | $0.897 \pm 0.032$ | 0.725-0.925 | $0.848 \pm 0.027$ | 0.714-0.909 | 2.2 |
| ARIBA | 0.612 ± 0.026 | 0.549-0.707 | 0.721 ± 0.018 | 0.639-0.763 | 0.533 ± 0.106 | 0.295-0.711 | 4.2 |
| KARGA | 0.549 ± 0.045 | 0.337-0.584 | 0.469 ± 0.065 | 0.272-0.588 | 0.695 ± 0.139 | 0.273-0.867 | 8.3 |
| SRST2 | 0.293 ± 0.063 | 0.055-0.373 | 0.213 ± 0.064 | 0.044-0.326 | 0.480 ± 0.158 | 0.072-0.691 | 21.5 |
| GROOT | 0.286 ± 0.085 | 0.067-0.396 | 0.267 ± 0.116 | 0.087-0.453 | 0.313 ± 0.086 | 0.051-0.452 | 29.8 |

Performance evaluation of five AMR detection tools across these datasets revealed distinct profiles (Figure 4b, Table 2). ARGprofiler achieved the highest mean F1-score (0.872 ± 0.019), along with high precision (0.897 ± 0.032) and recall (0.848 ± 0.027), and exhibited the lowest coefficient of variation (CV = 2.2%), indicating consistent performance. ARIBA and KARGA showed some variability, with CVs of 4.2% and 8.3%, respectively. In contrast, SRST2 and GROOT demonstrated substantially lower mean F1-scores (0.293 ± 0.063 and 0.286 ± 0.085) and higher variability (CVs of 21.5% and 29.8%, respectively).

Spearman’s rank correlation (ρ) between F1-score and the number of genomes per dataset was calculated (Table S2) to quantify the influence of community complexity on ARG detection. The correlation revealed tool-specific distinct tends: GROOT (ρ = 0.94, p < 4.75×10^-45^), KARGA (ρ = 0.76, p < 3×10^-20^), and SRST2 (ρ = 0.70, p < 0.001) showed strong positive associations with increasing complexity, whereas ARIBA (ρ = -0.59, p < 1.38×10^-10^) and ARGprofiler (ρ = -0.59, p < 9.83×10^-11^) exhibited moderate negative associations.

A Friedman rank-sum test revealed significant differences in F1-scores among the tools (χ^2^ = 379.28, *df* = 4, *p* < 2.2 × 10^−16^). Pairwise comparisons using the Nemenyi post-hoc test (single-step adjusted *p*-values) indicated that most tool pairs differed significantly (*p* < 0.05), except for GROOT and SRST2 (*p* = 0.9989), which performed comparably. Significant differences were observed between ARGprofiler and ARIBA (*p* = 6.2 × 10^−5^), GROOT and ARGprofiler (*p* < 2 × 10^−16^), GROOT and ARIBA (*p* = 7.8 × 10^−15^), KARGA and ARGprofiler (*p* = 3.5 × 10^−14^), KARGA and ARIBA (*p* = 1.1 × 10^−4^), KARGA and GROOT (*p* = 5.7 × 10^−11^), and SRST2 with all tools except GROOT (*p* < 0.01). The relative ranking of the tools based on their mean F1-scores is ARGprofiler > ARIBA > KARGA > SRST2 ≈ GROOT (Figure 4b).

### d. Effect of Log-Normal Coverage

The performance of the same set of tools was further evaluated on simulated metagenomic datasets with a lognormal distribution of sequencing coverage to mimic realistic metagenomic conditions. Performance was evaluated based on the detection of AMR genes, with ABRICATE-based predictions serving as the reference, and assessed using recall, precision, F1-score and CV (Table 3, Figure 4c). KARGA achieved highest mean F1-score (0.122 ± 0.067), supported by high precision (0.849 ± 0.135) and highest recall (0.067 ± 0.041). ARGprofiler (F1-score: 0.101 ± 0.075) exhibited highest precision overall (0.931 ± 0.095) and lower recall (0.55± 0.046) compared to KARGA. ARIBA followed with an F1-score of 0.086 ± 0.05, combining relatively high precision (0.618 ± 0.136) with low recall (0.048 ± 0.030). GROOT showed an F1-score of 0.050 ± 0.030, characterized by low precision (0.270 ± 0.196) and recall (0.030 ± 0.020). SRST2 exhibited the lowest performance (F1 = 0.002 ± 0.006) with minimal recall (0.001 ± 0.003) and poor precision (0.037 ± 0.150).

**Table 3:** Comprehensive Tool Performance Metrics Across 100 Datasets *CV = Coefficient of Variation (SD/Mean × 100)

| Tool | F1-Score Mean ± SD | F1-Score Range | Precision Mean ± SD | Precision Range | Recall Mean ± SD | Recall Range | F1-Score CV (%) |
| --- | --- | --- | --- | --- | --- | --- | --- |
| KARGA | 0.122 ± 0.067 | 0.028-0.351 | 0.849 ± 0.135 | 0.339-1.000 | 0.067 ± 0.041 | 0.014-0.232 | 54.78 |
| ARGprofiler | 0.101 ± 0.075 | 0.006-0.451 | 0.931 ± 0.095 | 0.500-1.000 | 0.055 ± 0.046 | 0.003-0.296 | 74.35 |
| ARIBA | 0.086 ± 0.051 | 0.011-0.248 | 0.618 ± 0.136 | 0.154-1.000 | 0.048 ± 0.030 | 0.006-0.149 | 58.72 |
| GROOT | 0.050 ± 0.030 | 0.000-0.128 | 0.270 ± 0.196 | 0.000-1.000 | 0.030 ± 0.020 | 0.000-0.093 | 59.83 |
| SRST2 | 0.002 ± 0.006 | 0.000-0.049 | 0.037 ± 0.150 | 0.000-1.000 | 0.001 ± 0.003 | 0.000-0.026 | 327.82 |

A Friedman rank-sum test confirmed significant differences in F1-scores among the tools (χ^2^ = 295.97, *df* = 4, *p* < 2.2 × 10^−16^). Subsequent pairwise comparisons using the Nemenyi post-hoc test (single-step adjusted *p-values*) revealed that all tool pairs differed significantly (*p* < 0.05) except ARGprofiler and ARIBA (*p* = 0.39). In particular, GROOT differed from ARGprofiler (*p* = 3.6 × 10^−10^) and ARIBA (*p* = 1.2 × 10−), KARGA differed from ARGprofiler (*p* = 0.02), ARIBA (*p* = 1.5 × 10^−5^), and GROOT (*p* = 3.6 × 10^−14^), while SRST2 differed significantly from all other tools (*p* < 10^−7^). The relative ranking of the tools based on their mean F1-scores showed that KARGA achieved the highest rank, followed by ARGprofiler, ARIBA, GROOT, and SRST2 (Figure 4c).

### e. Runtime and resource usage

We evaluated the real-time execution durations of ARGprofiler (KMA module), GROOT, KARGA, SRST2, and ARIBA to process 26Gb of data (i.e. 21 different coverage dataset), under standardised computational conditions using 16 CPUs and 100 GB of RAM.

The distribution of processing time of each tool is presented in Figure 5D. KARGA exhibited median processing time around 11 minutes (min) and values extending beyond ∼19 min. In contrast, KMA (median: ∼0.5 min) and SRST2 (median: ∼1.3 min) demonstrated the shortest and most consistent execution times. GROOT also performed efficiently, with processing time comparable to SRST2, albeit with slightly greater variability. ARIBA had a median processing time of approximately ∼6 minutes.

**Figure 5:**
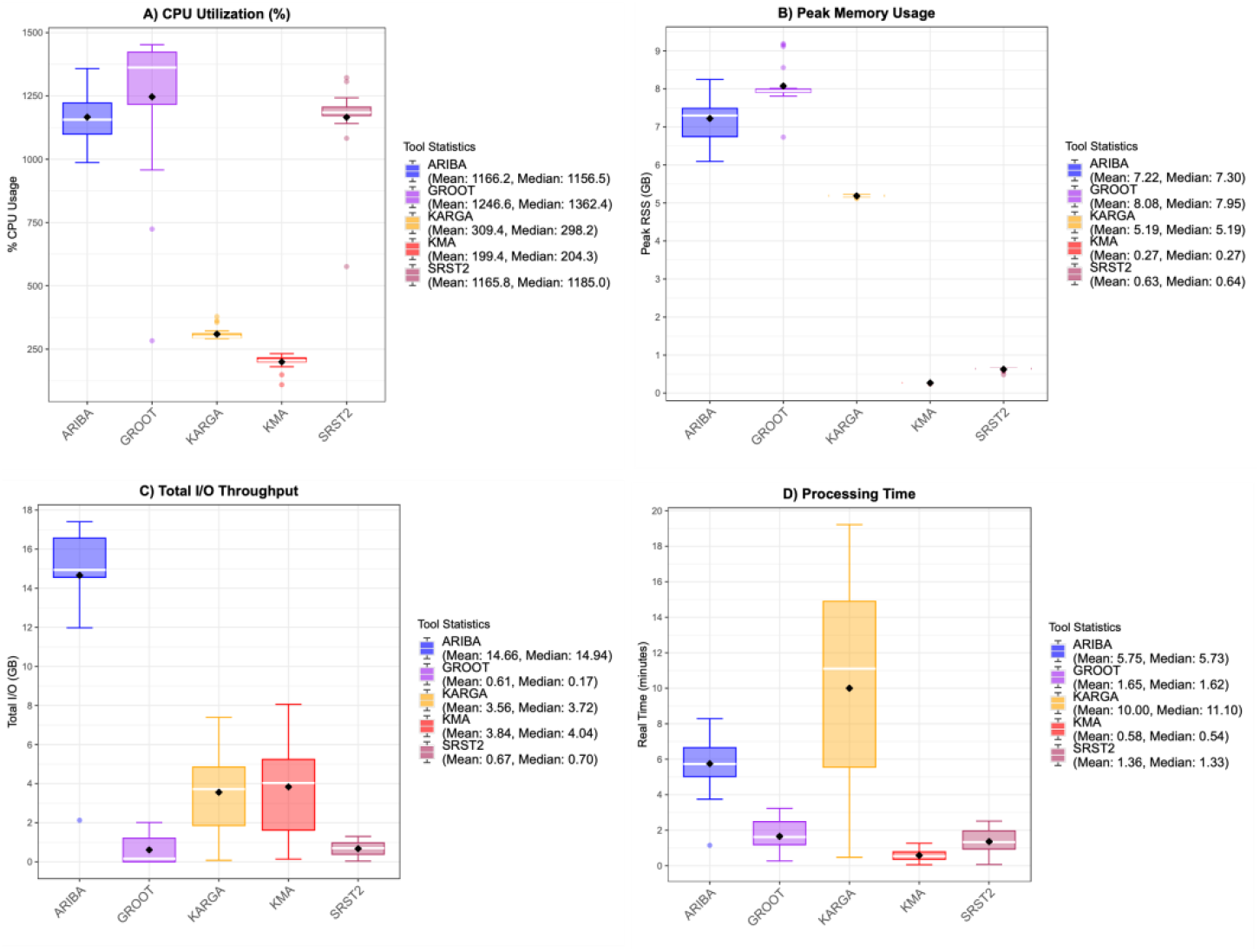
Performance comparison of five tools for ARGs detection on variable sequencing depth (1x-100x dataset). Box plots showing (A) CPU utilization, (B) Peak memory usage (RSS: Resident Set Size), (C) Total I/O throughput, and (D) Processing time across 21 samples processed by each tool. Each box represents the interquartile range (IQR) with the white line indicating the median and the black diamond indicating the mean. Whiskers extend to 1.5 × IQR from the quartiles, with outliers shown as open circles. Tools are color-coded as follows: ARIBA (blue), GROOT (purple), KARGA (orange), KMA (red), and SRST2 (maroon). The legend for each panel displays the median and mean values for each tool. All tools were allocated 16 CPUs and run on identical input data. CPU utilization exceeding 100% indicates effective multi-threading.

CPU utilization patterns (Figure 5A) revealed substantial differences in parallelisation efficiency. GROOT achieved the highest median CPU utilization at 1,362% followed by ARIBA (1,157%) and SRST2 (1,185%), respectively. KMA and KARGA showed lower CPU utilisation (median: 204% and 298%, respectively).

Maximum memory consumption (Figure 5B) varied considerably across tools. GROOT required the highest memory with a median of 7.95 GB, followed by ARIBA (7.30 GB). KARGA maintained a consistent memory footprint around ∼5.2 GB. SRST2 used substantially less memory (median: 652 MB), while KMA was the most memory-efficient at 278 MB.

I/O throughput analysis (Figure 5C) showed that ARIBA generated the highest data transfer volumes with a median total I/O of 14.94 GB, followed by KMA and KARGA I/O activity (∼4.0 GB and ∼4.3 GB, respectively), while SRST2 and GROOT had the lowest I/O footprints (719 MB and 169 MB, respectively).

Overall, KMA, SRST2, and GROOT were the most time-efficient tools, while ARIBA provided a balance between computational efficiency and analytic depth. KARGA was the most resource-intensive, which may be an important consideration depending on dataset size, pipeline throughput requirements, or available infrastructure.

## 5. Discussion

In this study, we systematically evaluated the impact of sequencing coverage and microbial complexity on the detection of antimicrobial resistance genes (ARGs) from shotgun metagenomic data. By employing a comprehensive and unified ARG reference database. We excluded ARGs that require SNPs to confer resistance from our analysis, thereby restricting the analysis to acquired ARGs detectable at gene level. We evaluated read-based detection methods, selected according to criteria outlined earlier. To disentangle the effects of sequencing coverage and community composition, these two factors were varied independently. Sequencing coverage was systematically altered while keeping the community composition constant to assess the effect of coverage.Conversely, to evaluate the effect of community complexity, additional genomes were systematically added to mock communities while maintaining a constant sequencing coverage. Finally, we assess the performance of these tools on approximate realistic metagenomic conditions, where raw reads were simulated with lognormal coverage distributions. A summary of tool performance across all experimental conditions is provided in table 4.

**Table 4:** The tools performance was summarized in this table. Colours are scaled from minimum to maximum value for each experimental condition. The tools are ordered based on high to low based on their mean F1-score in each experimental condition.

| Tools | Time (in Min) | Recall | Precision | F1-score | Experimental Condition |
| --- | --- | --- | --- | --- | --- |
| ARGprofiler | 0.542 | 0.805 | 0.861 | 0.815 | Variable Coverage, Fixed Community |
| ARIBA | 5.73 | 0.729 | 0.754 | 0.724 | Variable Coverage, Fixed Community |
| KARGA | 11.1 | 0.527 | 0.587 | 0.522 | Variable Coverage, Fixed Community |
| GROOT | 1.62 | 0.315 | 0.176 | 0.19 | Variable Coverage, Fixed Community |
| SRST2 | 1.33 | 0.147 | 0.267 | 0.182 | Variable Coverage, Fixed Community |
| ARGprofiler | 9.53 | 0.848 | 0.897 | 0.872 | Fixed Coverage, Increased Community |
| ARIBA | 37.3 | 0.533 | 0.721 | 0.612 | Fixed Coverage, Increased Community |
| KARGA | 144 | 0.695 | 0.469 | 0.549 | Fixed Coverage, Increased Community |
| GROOT | 19.6 | 0.48 | 0.213 | 0.293 | Fixed Coverage, Increased Community |
| SRST2 | 17.6 | 0.313 | 0.267 | 0.286 | Fixed Coverage, Increased Community |
| KARGA | 2.28 | 0.067 | 0.849 | 0.122 | Approximate Realistic Community |
| ARGprofiler | 0.126 | 0.055 | 0.931 | 0.101 | Approximate Realistic Community |
| ARIBA | 8.48 | 0.048 | 0.618 | 0.086 | Approximate Realistic Community |
| GROOT | 0.633 | 0.03 | 0.27 | 0.05 | Approximate Realistic Community |
| SRST2 | 0.378 | 0.001 | 0.037 | 0.002 | Approximate Realistic Community |

The integration of seven publicly available ARG databases yielded a comprehensive and non-redundant gene collection. Despite the substantial overlap among existing databases, the pairwise comparisons revealed the pronounced heterogeneity in gene content. Approximately 80% of the genes were shared among the databases, yet none were common to all. The mean pairwise Jaccard similarity was low, with highest concordance between AMRFinderPlus and MEGARes, while (encompassing metal resistance genes) and ResFinderFG (containing metagenome-derived ARGs) exhibited minimal overlap with other databases. Together, these findings highlight the inconsistency in ARG database composition and suggest that a single database may not capture the full resistome diversity. Therefore, using multiple databases, either through integrating them into a unified reference set or by harmonizing outputs from tools that use different databases ^42^, can enhance the comprehensiveness of ARGs profiling in metagenomic studies. However, harmonizing gene resources remains difficult because, although genes are often grouped by function or sequence similarity, databases differ in annotation standards, nomenclature conventions, and ontological frameworks. Hence, making it challenging to derive universally accepted gene names or functional annotations.

Our findings demonstrated significant differences in the performance of the five tools across sequencing coverages when evaluated on a fixed microbial community. A minimum coverage of 10x was required for reliable detection of ARGs, as lower depth reduce recall. Detection performance stabilized beyond 10x coverage, with only minor improvement observed beyond 30x. Optimal detection performance is achieved between 20×–30×, where precision and recall are maximized, and improvements begin to plateau beyond this range. Among the evaluated tools, ArgProfiler consistently outperformed others, achieving a mean F1-score of 0.891 at ≥10× coverage. ARIBA also performed well (mean F1= 0.778 ± 0.020 at ≥10×). KARGA showed moderate overall performance and relatively higher recall at lower coverage (<10×), and its precision was near to ARGprofiler at same coverage (<10×). In contrast, GROOT and SRST2 performed poorly (mean F1 < 0.21) under identical conditions primarily due to low recall. These results confirm that sequencing coverage significantly influences ARGs detection, particularly in metagenomic datasets. Tools originally designed for whole-genome sequencing (WGS) of isolates, such as SRST2, are unsuitable for metagenomic applications and should be used with caution. ARIBA, although developed for isolate-level analyses, showed moderate and consistent performance at sufficient coverage, suggesting conditional applicability in metagenomic contexts.

Building on this, we next evaluated how the increasing microbial community complexity affects detection when sequencing coverage was fixed (at 25×). Across 100 systematically constructed mock communities of increasing genomic diversity, all tools exhibited distinct performance. ARGprofiler maintained the highest and most consistent performance (mean F1 = 0.872 ± 0.019; CV = 2.2%). ARIBA and KARGA showed moderate variability, while SRST2 and GROOT displayed markedly lower F1-scores and higher variability. However, correlation analysis revealed contrasting trends. GROOT, KARGA, and SRST2 exhibited strong positive correlations between F1-score and community size, indicating improved detection as complexity increased, whereas ARGprofiler and ARIBA showed moderate negative correlations, suggesting a slight decline in accuracy under highly diverse conditions. Statistical comparisons confirmed significant performance differences among tools, with ARGprofiler and ARIBA outperforming the others overall.

Finally, to better approximate realistic metagenomic conditions, the tools were evaluated on 100 simulated datasets with a log-normal distribution of sequencing coverage, reflecting the uneven read coverage typically observed in metagenomic samples. Under these conditions, all tools exhibited a pronounced decline in recall and exhibited higher variability of performance across the datasets, than earlier described experiment settings. Overall, the mean F1-score is quite low as compared to either variable coverage (with constant community) or f increased complexity (with constant coverage). Statistical analysis confirmed significant differences among all tools (p < 0.001), with KARGA achieving the highest mean F1-score (0.122 ± 0.067), supported by high precision, followed by ARGprofiler and ARIBA. Consistent with previous observations, GROOT and SRST2 demonstrated poor performance, with mean F1-score below 0.05.

Collectively, these findings demonstrate that sequencing coverage and community complexity jointly shape the performance of ARG detection tools. While ARGprofiler and ARIBA performed relatively well under uniform coverage, their sensitivity decreased in unevenly distributed and highly complex communities. In contrast, KARGA showed greater adaptability under realistic conditions. Additionally, GROOT demonstrated consistently lower recall through, although developed for predicating ARGs from metagenomic datasets. Hence, the choice of the algorithm to detect ARGs in metagenome is critically important and it does rely on complexity and sequencing coverage of the metagenome sample.

In addition to detection accuracy, computational efficiency plays a critical role in determining tool suitability for large-scale or time-sensitive analyses. Our benchmarking analysis revealed substantial differences in runtime and resource consumption. ARGprofiler, SRST2, and GROOT were the most time-efficient, offering rapid execution with minimal memory usage. Among them, ARGprofiler demonstrated the lowest CPU and memory requirements, making it particularly suitable for high-throughput or resource-limited environments. GROOT, while slightly more memory-intensive, benefited from efficient parallelization and low I/O overhead. ARIBA balanced moderate computational demands with analytic depth, though its higher I/O load may affect scalability. In contrast, KARGA, despite its competitive accuracy, imposed the heaviest computational burden across all metrics, limiting its practicality for large datasets or real-time diagnostic pipelines without substantial computing resources.

However, this study as several limitations that should be considered when interpreting the findings. Although the simulated datasets were systematically designed to emulate metagenomic conditions, they may not fully capture the biological and technical variability present in real microbiomes, including uneven taxonomic abundances, and sequencing biases. In addition, the use of reference ARG database helps to screen for only well characterized ARGs not for novel or poorly characterized ARGs. Furthermore, ABRICATE, which uses assembly to predict ARGs within them may itself constrained by the quality of assemblies, potentially missing, fragmented or low-abundance ARGs. Nevertheless, if a gene is detected on assembly at 90% identity and 90% coverage, there is a high chance that the gene is present in the sample. The evaluation also did not assess the performance across different resistance classes, such as such as β-lactamases or efflux pumps. Overall, these factors highlight the need for continued benchmarking using real metagenomic datasets and diverse resistance determinants to better capture the complexity of ARG detection in natural microbial communities. Finally, these remains a pressing need for high-quality, standardized metagenomic datasets that realistically represent microbial community complexity and ARGs abundance, which can serve as ground-truth resources for future benchmarking efforts.

## Note

Portions of this manuscript were refined with the assistance of large language models, including ChatGPT (GPT-5, OpenAI, 2025) and Claude (Sonnet 4.5, Anthropic, 2025), which were used exclusively for language polishing and paraphrasing. These models were also employed to generate code for data visualization; all code outputs were manually reviewed, tested, and verified for correctness using example datasets. The models were not involved in data analysis, interpretation, or generation of scientific results. All analyses and conclusions presented in this study were performed independently by the authors, who take sole responsibility for the integrity and accuracy of the work.

## Acknowledgements

The authors gratefully acknowledge the support of the Biotechnology and Biological Sciences Research Council (BBSRC); this research was funded by the BBSRC Institute Strategic Programme Food Microbiome and Health BB/X011054/1 and its constituent project(s) BBS/E/QU/230001B; the BBSRC Institute Strategic Programme Microbes and Food Safety BB/X011011/1 and its constituent project(s) BBS/E/QU/230002C; the BBSRC Core Capability Grant BB/CCG2260/1. Computing infrastructure was in part provided by MRC CLIMB BIG DATA grant MR/T030062/1 and in part by NBI Research Computing.

## Authors’ contributions

**AP, AT, and SKT** conceptualized the work. **AP and SKT** design the benchmarking framework. **SH and SKT** building NextFlow pipeline. **JT and SKT** result analysis and visualization. **SKT** produced results and wrote the manuscript. **AP, AT, KG, JT, and SH** contributed to interpretation and revision of manuscript. **AP, AT, KG, JT, SH and SKT** read and approved the final manuscript.

## Data availability statement

The pipeline is available here: https://github.com/quadram-institute-bioscience/ARG-Sniper; The panARG database is available 10.5281/zenodo.17611955;Custom reference databases: 10.5281/zenodo.18429574

## Notes

https://github.com/quadram-institute-bioscience/ARG-Sniper

### Competing Interest Statement

The authors have declared no competing interest.

## Reference

1. Murray, C. J. L. et al. Global burden of bacterial antimicrobial resistance in 2019: a systematic analysis. The Lancet 399, 629–655 (2022).

2. Naghavi, M. et al. Global burden of bacterial antimicrobial resistance 1990–2021: a systematic analysis with forecasts to 2050. The Lancet 404, 1199–1226 (2024).

3. Beghini, F. et al. Integrating taxonomic, functional, and strain-level profiling of diverse microbial communities with bioBakery 3. Elife 10, e65088 (2021).

4. Lema, N. K., Gemeda, M. T. & Woldesemayat, A. A. Recent Advances in Metagenomic Approaches, Applications, and Challenges. Curr Microbiol 80, 347 (2023).

5. Wood, D. E., Lu, J. & Langmead, B. Improved metagenomic analysis with Kraken 2. Genome Biol 20, 257 (2019).

6. Christian, B. et al. Long-Term Temporal Stability of the Resistome in Sewage from Copenhagen. mSystems 5, 10.1128/msystems.00841-20 (2020).

7. Martiny, H.-M., Munk, P., Brinch, C., Aarestrup, F. M. & Petersen, T. N. A curated data resource of 214K metagenomes for characterization of the global antimicrobial resistome. PLoS Biol 20, e3001792. (2022).

8. de Abreu, V. A. C., Perdigão, J. & Almeida, S. Metagenomic Approaches to Analyze Antimicrobial Resistance: An Overview. Front Genet Volume 11-2020, (2021).

9. Abramova, A., Karkman, A. & Bengtsson-Palme, J. Metagenomic assemblies tend to break around antibiotic resistance genes. BMC Genomics 25, 959 (2024).

10. Rowe, W. P. M. & Winn, M. D. Indexed variation graphs for efficient and accurate resistome profiling. Bioinformatics 34, 3601–3608 (2018).

11. Prosperi, M. & Marini, S. KARGA: Multi-platform Toolkit for k-mer-based Antibiotic Resistance Gene Analysis of High-throughput Sequencing Data. in 2021 IEEE EMBS International Conference on Biomedical and Health Informatics (BHI) 1–4 (IEEE, 2021). doi:10.1109/BHI50953.2021.9508479.

12. Martiny, H.-M. et al. ARGprofiler—a pipeline for large-scale analysis of antimicrobial resistance genes and their flanking regions in metagenomic datasets. Bioinformatics 40, (2024).

13. Seemann T. ABRICATE. Preprint at https://github.com/tseemann/abricate (2015).

14. Feldgarden, M. et al. AMRFinderPlus and the Reference Gene Catalog facilitate examination of the genomic links among antimicrobial resistance, stress response, and virulence. Sci Rep 11, 12728 (2021).

15. Berglund, F. et al. Identification and reconstruction of novel antibiotic resistance genes from metagenomes. Microbiome 7, 52 (2019).

16. Lakin, S. M. et al. Hierarchical Hidden Markov models enable accurate and diverse detection of antimicrobial resistance sequences. Commun Biol 2, 294 (2019).

17. Arango-Argoty, G. et al. DeepARG: a deep learning approach for predicting antibiotic resistance genes from metagenomic data. Microbiome 6, 23 (2018).

18. Pei, Y. et al. ARGNet: using deep neural networks for robust identification and classification of antibiotic resistance genes from sequences. Microbiome 12, 84 (2024).

19. Li, Y. et al. HMD-ARG: hierarchical multi-task deep learning for annotating antibiotic resistance genes. Microbiome 9, 40 (2021).

20. Jia, B. et al. CARD 2017: expansion and model-centric curation of the comprehensive antibiotic resistance database. Nucleic Acids Res 45, D566–D573 (2017).

21. Alcock, B. P. et al. CARD 2023: expanded curation, support for machine learning, and resistome prediction at the Comprehensive Antibiotic Resistance Database. Nucleic Acids Res 51, D690–D699 (2023).

22. Florensa, A. F., Kaas, R. S., Clausen, P. T. L. C., Aytan-Aktug, D. & Aarestrup, F. M. ResFinder – an open online resource for identification of antimicrobial resistance genes in next-generation sequencing data and prediction of phenotypes from genotypes. Microb Genom 8, 000748 (2022).

23. Pal, C., Bengtsson-Palme, J., Rensing, C., Kristiansson, E. & Larsson, D. G. J. BacMet: antibacterial biocide and metal resistance genes database. Nucleic Acids Res 42, D737–D743 (2014).

24. Bonin, N. et al. MEGARes and AMR++, v3.0: an updated comprehensive database of antimicrobial resistance determinants and an improved software pipeline for classification using high-throughput sequencing. Nucleic Acids Res 51, D744–D752 (2023).

25. Gschwind, R. et al. ResFinderFG v2.0: a database of antibiotic resistance genes obtained by functional metagenomics. Nucleic Acids Res 51, W493–W500 (2023).

26. Wissel, E. F. et al. hAMRoaster: a tool for comparing performance of AMR gene detection software. bioRxiv 2022.01.13.476279 (2023) doi:10.1101/2022.01.13.476279.

27. Rooney, A. M. et al. Performance Characteristics of Next-Generation Sequencing for the Detection of Antimicrobial Resistance Determinants in Escherichia coli Genomes and Metagenomes. mSystems 7, (2022).

28. Kurtzer, G. M., Sochat, V. & Bauer, M. W. Singularity: Scientific containers for mobility of compute. PLoS One 12, e0177459. (2017).

29. Di Tommaso, P. et al. Nextflow enables reproducible computational workflows. Nat Biotechnol 35, 316–319 (2017).

30. Baker-Austin, C., Wright, M. S., Stepanauskas, R. & McArthur, J. V. Co-selection of antibiotic and metal resistance. Trends Microbiol 14, 176–182 (2006).

31. Kumar, G. S. et al. ARG-ANNOT, a New Bioinformatic Tool To Discover Antibiotic Resistance Genes in Bacterial Genomes. Antimicrob Agents Chemother 58, 212–220 (2014).

32. Pal, C., Bengtsson-Palme, J., Rensing, C., Kristiansson, E. & Larsson, D. G. J. BacMet: antibacterial biocide and metal resistance genes database. Nucleic Acids Res 42, D737–D743 (2014).

33. Martiny, H.-M. et al. ARGprofiler—a pipeline for large-scale analysis of antimicrobial resistance genes and their flanking regions in metagenomic datasets. Bioinformatics 40, btae086 (2024).

34. Hunt, M. et al. ARIBA: Rapid antimicrobial resistance genotyping directly from sequencing reads. Microb Genom 3, (2017).

35. Edgar, R. C. Search and clustering orders of magnitude faster than BLAST. Bioinformatics 26, 2460–2461 (2010).

36. Hiseni, P., Rudi, K., Wilson, R. C., Hegge, F. T. & Snipen, L. HumGut: a comprehensive human gut prokaryotic genomes collection filtered by metagenome data. Microbiome 9, 165 (2021).

37. Gourlé, H., Karlsson-Lindsjö, O., Hayer, J. & Bongcam-Rudloff, E. Simulating Illumina metagenomic data with InSilicoSeq. Bioinformatics 35, 521–522 (2019).

38. Olm, M. R., Brown, C. T., Brooks, B. & Banfield, J. F. dRep: a tool for fast and accurate genomic comparisons that enables improved genome recovery from metagenomes through de-replication. ISME J 11, 2864–2868 (2017).

39. Rowe, W. P. M. & Winn, M. D. Indexed variation graphs for efficient and accurate resistome profiling. Bioinformatics 34, 3601–3608 (2018).

40. Inouye, M. et al. SRST2: Rapid genomic surveillance for public health and hospital microbiology labs. Genome Med 6, 90 (2014).

41. Prosperi, M. & Marini, S. KARGA: Multi-platform Toolkit for k-mer-based Antibiotic Resistance Gene Analysis of High-throughput Sequencing Data. in 2021 IEEE EMBS International Conference on Biomedical and Health Informatics (BHI) 1–4 (2021). doi:10.1109/BHI50953.2021.9508479.

42. Mendes, I. et al. hAMRonization: Enhancing antimicrobial resistance prediction using the PHA4GE AMR detection specification and tooling. Preprint at 10.1101/2024.03.07.583950 (2024).

